# Endotoxin Tolerance Induced by Different TLR Ligands

**DOI:** 10.1101/250415

**Authors:** Suzanne K. Butcher, Christine O’Carroll, Christine A. Wells, Ruaidhrí J. Carmody

## Abstract

Endotoxin tolerance is a long-recognised property of macrophages that leads to an altered response to repeated or chronic exposure to Toll-like receptor (TLR) ligands. The physiological role of tolerance is to limit the potential damage to host tissue that may otherwise result from prolonged production of pro-inflammatory cytokines. Endotoxin tolerance is induced by all TLRs tested to date, however, tolerance induced by the TLR4 ligand lipopolysaccharide (LPS) is by far the best studied. LPS tolerance involves a global transcriptional shift from a pro-inflammatory response toward one characterised by the expression of anti-inflammatory and pro-resolution factors. Although largely reversible, LPS-tolerance leads to a hybrid macrophage activation state that is pro-inflammatory in nature but possesses distinct regulatory anti-inflammatory features. Remarkably, a comparative transcriptomic analysis of tolerance induced by different TLR ligands has not previously been reported. Here we describe the transcriptomic profiles of mouse macrophages tolerised with ligands for TLR2, TLR3, TLR4 and TLR 9. While we identified TLR-specific transcriptional profiles in macrophages tolerised with each ligand, tolerance induced by TLR4 was the most comprehensive state, such that each gene tolerised by any of the TLRs tested was also found to be tolerised by TLR4. Pro-inflammatory cytokines are not universally supressed in all tolerant cells but distinct patterns of cytokine expression distinguished TLR-specific tolerance. Analysis of gene regulatory regions revealed specific DNA sequence motifs associated with distinct states of TLR tolerance, implicating previously identified as well as novel transcriptional regulators of tolerance in macrophages. These data provide a basis for the future exploitation of TLR-specific tolerant states to achieve therapeutic re-programming of the innate immune response.

## Introduction

Innate immunity is the first line of host defence against infection and is critical for the development of adaptive immunity. Toll-like receptors (TLRs) are key sensors in the innate immune system and recognise conserved structures of microbial-derived molecules or pathogen associated molecular patterns (PAMPs). To date, 13 TLRs have been identified in mammals. These form homo‐ or heterodimers to recognise a range of PAMPs that spans microbial diversity, and regulation of this TLR repertoire fundamentally alters the tissue response to infection (reviewed in (Huang and Wells, 2014; Leifer and Medvedev, 2016)). TLR activation induces the expression of hundreds of genes that encode inflammatory cytokines, type I interferons, antimicrobial proteins, and regulators of metabolism and regeneration; these molecules in turn mediate inflammation, anti-microbial immunity and tissue regeneration.

The activation of different TLRs leads to specific transcriptional responses, *via* highly evolutionarily conserved signalling pathways. These are dependent on the adapter proteins MyD88, TRIF, TIRAP and TRAM, which direct activation of the NF-κB, MAPK and IRF pathways (McGettrick and O’Neill, 2010). The combination of adapter proteins engaged by specific TLRs shapes the subsequent transcriptional and immune responses to the initiating ligand. The choice of adapter can be mediated by subcellular location of the TLR-pathogen engagement, with intracellular (endosomal) TLRs preferentially signalling via non-MyD88 pathways. The synergistic activation of NF-κB, MAPK and IRF pathways are important for activation of acute cytokine responses, including TNF, IL6 and IL1β.

The negative regulation of TLR-signalling events is critical to ensure that prolonged or repeated exposure to TLR ligands does not lead to uncontrolled or inappropriate inflammation and consequent damage to host tissue. The most important mechanism for controlling TLR activation is a form of tolerance to repeated exposure to a TLR ligand. This has been best described for lipopolysaccharide (LPS) activation of TLR4 and is otherwise known as endotoxin-tolerance (Medvedev, Kopydlowski and Vogel, 2000; Collins and Carmody, 2015). TLR-tolerance can be described as a state of altered responsiveness of cells to the repeated or chronic activation of TLRs, and includes the phenomena of cross-tolerance, where pre-exposure to one TLR-ligand will reduce inflammatory responses to another (Dobrovolskaia *et al.*, 2003; Julian *et al.*, 2015). While convergent signalling *via* NF-κB is essential for acquisition of LPS-tolerance, it is not known how generalizable this may be to other TLR-ligands (Dobrovolskaia *et al.*, 2003; Carmody *et al.*, 2007). Chromatin changes at tolerised genes allow for persistence of altered responsiveness to re-stimulation of cells, but these changes are reversible over time, or in response to competing signals (Foster, Hargreaves and Medzhitov, 2007; Novakovic *et al.*, 2016).

Previous transcriptomic analysis of LPS (TLR4) tolerant cells identified two classes of TLR4-inducible genes; i) tolerisable genes which are repressed during LPS tolerance, and ii) non-tolerisable genes, which are not (Foster, Hargreaves and Medzhitov, 2007; Mages, Dietrich and Lang, 2008; O’Carroll *et al.*, 2014). The functional classification of LPS-inducible genes revealed that pro-inflammatory factors fall predominantly into the tolerisable class of genes, while genes which code for anti-microbial factors, including anti-microbial peptides and scavenger receptors, fall into the non-tolerisable class of genes. Thus, LPS tolerance represents a global transcriptional shift from a pro-inflammatory to a pro-resolution and anti-inflammatory response, while maintaining protective innate immune functionality in the context of chronic or continuing infection. Whether genes are tolerised or not likely reflects the impact of continued expression in the context of an inflammatory response and whether repression would be advantageous or deleterious. Furthermore, LPS tolerance is a transient state that allows cells to re-express pro-inflammatory factors in response to TLR ligands over time. Our previous transcriptomic analysis demonstrated that macrophages that have recovered from a tolerant state adopt a hybrid polarisation state with features of both M1 and M2 macrophages (O’Carroll *et al.*, 2014).

To date, global transcriptomic analysis has only been performed for tolerance induced by LPS, and the similarity to tolerance induced by ligands for other TLRs is not known. In this study, we perform a comparative transcriptomic analysis of murine bone marrow derived macrophages (BMDM) tolerised with ligands for TLR2, TLR3, TLR4 and TLR9. Our analysis identifies a core set of genes tolerised by all TLR ligands tested, and further reveals a pattern of LPS‐ TLR4 tolerance that exemplifies tolerance in TLR2, TLR3 and TLR9. We identified additional patterns of super-repression and super-induction on re-stimulation that indicate a diverse set of transcriptional events that shape the long-term response of macrophages to infection.

## Materials and methods

### Murine bone marrow isolation

Bone marrow was isolated from 6-8 weeks old female C57BL/6 for generation of primary bone marrow derived macrophages (BMDM) *in vitro*. Mice were sacrificed according to the Code of Practice for the Humane Killing of Animals under Schedule 1 to the UK Animals (Scientific Procedures) Act 1986. Excess tissue was removed from the femur and tibia bones and then cleaned in sterile phosphate buffered saline (PBS) and 70% ethanol. Using a 21-gauge needle and syringe, bone marrow was isolated by flushing ice cold sterile PBS through the femur and tibia bones. Isolated bone marrow was re-suspended to generate a single cell suspension and passed through a 70μM cell strainer to remove any debris. The bone marrow suspension was washed twice in culture media (DMEM, 10% FBS, 1% penicillin/streptomycin, 1% LGlutamine, 1% non-essential amino acids), and centrifuged at 4°C at 300 × g for 5 minutes. Bone marrow was cryopreserved in foetal calf serum (FCS) supplemented with 10% DMSO until required for use.

### Bone marrow derived macrophage differentiation

Bone marrow was cultured following isolation or from cryopreserved stocks in culture media supplemented with 30% L929 conditioned media for seven days. The cells were cultured on sterile non-tissue culture treated petri dishes. On day 3, BMDM differentiation media was removed and replaced with fresh differentiating BMDM culturing media, with any non-adherent cells removed at this stage. Adherent monocytes/macrophage progenitors differentiated into BMDMs by day 7. Differentiated BMDMs were removed from the petri dishes by incubating the cells with 5mM EDTA in sterile PBS at 37°C for 5 minutes. Cells were washed twice in BMDM culture media at 4°C for 5 minutes at 300 × g, re-suspended in BMDM culture media and transferred to tissue culture treated dishes for downstream experimental use. The purity of BMDMs was assessed by flow cytometry and was typically greater than 95% F4/80 positive.

### Endotoxin tolerance

Endotoxin tolerance was induced in BMDMs by stimulating cells for 24 hours with 100ng/ml LPS (Invivogen), 100ng/ml Pam3CSK4 (Invivogen), 10μg/ml Poly (I:C) (GE Healthcare), or 1μM CpG (1826 sequence, Eurofins). After 24 hours, the media was removed and the cells were washed twice with sterile PBS. The cells were allowed to rest in fresh culture media for 1 hour before a second stimulation for 4 hours with the same TLR ligand (Figure 1 (A)).

### Immunoblotting

Whole-cell proteins were extracted using RIPA lysis buffer supplemented with protease inhibitors (50 mM Tris•HCl, pH 7.4, 1% Nonidet P-40, 0.25% SDS, 150 mM NaCl, 1 mM EDTA, 1mM PMSF, 1mMNaF, 1mM Na3VO4, 2μg/ml aprotinin, 1μg/ml pepstatin, and 1μg/ml leupeptin). Lysates were resolved on SDS-PAGE gels, transferred to nitrocellulose membranes and immunoblotted with anti-p105/p50 (Cell Signaling Technology), anti-BCL-3 (AbCam).

### Gene expression analysis

Total RNA was isolated using the RNeasy mini kit (Qiagen) with all samples DNase treated according to manufacturer’s instructions. Triplicate biological replicate samples submitted for microarray profiling met all sample submission criteria (Beckman Coulter Genomics, NC, USA). Briefly, 200ng of total RNA was fluorescently labelled with Cy3 nucleotides. Labelled RNA (cRNA) was hybridised to Agilent mouse 8 × 60K microarrays (Agilent-028005). Each BMDM culture was generated from bone marrow pooled from three mice, and 2-3 biological replicates were performed per condition. Real-time PCR (qPCR) was performed using the universal probe library system (Roche) and primer sequences as follows; *Il6* forward 5’tctaattcatatcttcaaccaagagg3’ *Il6* reverse 5’-tggtccttagccactccttc-3; *Tnf* forward 5’-tcttctcattcctgcttgtgg-3’ *Tnf* reverse 5’-ggtctgggccatagaactga-3’; 18s forward 5’-aaatcagttatggttcctttggtc-3’ 18s reverse 5’-gctctagaattaccacagttatccaa-3’. Relative mRNA levels were calculated using the ΔΔCT method.

Microarray data was processed as follows: Data was normalised using Limma (3.26.9), including RMA background correction quantile normalisation (Ritchie *et al.*, 2015). Analysis workflow is described in Figure 1 (B): briefly, only probes mapping to an ENSEMBL (v67) gene were retained for this analysis. A detection threshold was applied to remove probes that were not expressed in at least 2/3 replicates. A linear model was fitted to identify differentially expressed probes using an adjusted p-value (Benjamini and Hochberg) of 0.05. A tolerised gene was defined by the following two criteria: where the same probe was (a) significantly induced (p < 0.05) and exhibited 1.5-fold or more inducible expression in response to the first stimulation, then (b) 1.5-fold lower induction in the re-stimulated condition. Tolerised probes were grouped by Partioning Around Medoids (PAM) clustering (R ‘cluster’ package (v2.0.6)) (Maechler *et al.*, 2017). Functional enrichment analysis was conducted on the top 500 differentially expressed genes in each condition, ranked but not filtered on p-value. Enriched pathways and GO terms were identified using the curated data at InnateDB (Breuer *et al.*, 2013), and protein-protein interactions were identified using STRINGDB (Franceschini *et al.*, 2013). Transcription factor motif enrichment was identified using Hypergeometric Optimization of Motif EnRichment (HOMER (v4.8.3)) (Heinz *et al.*, 2010). Heatmaps were generated using a similarity metric derived from the Pearson correlation, or using the www.stemformatics.org hierarchical clustering tool. Raw data is available from GEO (GSE81291) and www.stemformatics.org (Wells *et al.*, 2013) and processed data can be visualised at http://www.stemformatics.org/datasets/search?ds_id=6943.

**Figure 1:**
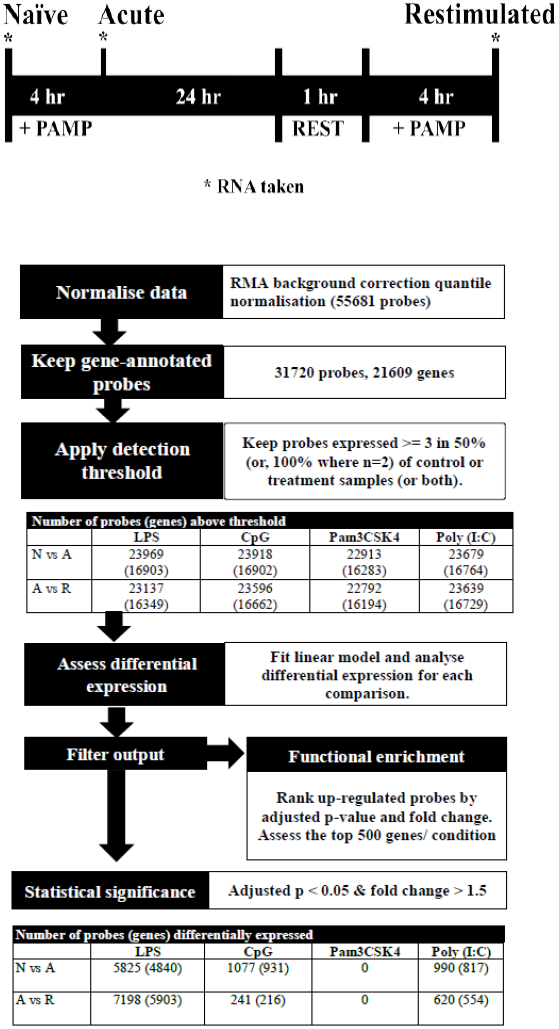
Experimental overview and workflow for transcriptome data normalisation, filtering and analysis. **(A)** Experimental procedure for induction of tolerance in mouse BMDMs. **(B)** Workflow for data normalisation, filtering and assessment of differential expression.

## Results

### A comparative analysis of TLR tolerance identifies TLR4 induced tolerance as the dominant form

LPS is classically used to study macrophage endotoxin tolerance. Although tolerance may be induced by TLRs other than TLR4, a comparative transcriptomic analysis of tolerance induced by different TLRs has not previously been performed. The similarity between transcriptional responses in macrophages tolerised by ligands for TLR2, TLR3, TLR4 and TLR9 following re-stimulation was assessed in mouse bone marrow derived macrophages (BMDM) stimulated with Pam3CSK4 (TLR2), Poly (I:C) (TLR3), LPS (TLR4) and CpG DNA (TLR9). 24 hours following stimulation the cells were washed then re-stimulated with the same TLR ligand for an additional 4 hours, prior to RNA isolation and microarray analysis.

We first confirmed that all four stimuli drove an acute activation profile by taking RNA at 4hr post-stimulation in BMDM that had received no prior stimulus. As shown in Figure 2, and accompanying Supplementary information (Figure S1, Table S1), we evaluated the top 500 BMDM genes activated by each ligand, to demonstrate a robust activation profile for known inflammatory markers, and a network of commonly activated genes that include chemokines and cytokines, and is centred on NF-κB, MAPK and IRF hubs. Assigning functional terms to the top 500 genes in each activation series (Supplementary Table S2) shows that although distinct patterns of gene expression are evident for each TLR ligand, these converge on similar biological processes.

Whereas NF-κB activation is a common theme to all the TLR ligands profiled here, the MAPK signalling pathways were more significantly enriched in the gene sets induced by CpG and Pam3CSK4, and type I interferon pathways were enriched in Poly (I:C) and LPS-driven gene sets. A set of 198 genes (Supplementary Table S1) up-regulated by LPS and Poly (I:C), but not CpG and Pam3CSK4, was rich in genes which regulate, and are driven by type I interferons. This included positive regulators *Azi2* and *Isg15*; and genes for IRF3/7-activating DNA sensors cGAS (*E330016A19Rik*), ZBP1, RIG-I (*Dhx58*), and MDA5 (*Ifih1*) Negative regulators of type I interferon signalling were also induced, including *Nlrc5* and *Usp18*. LPS and Poly (I:C) driven gene sets were significantly enriched for transcription factor motifs for IRF family members (Supplementary Figure S1).

**Figure 2:**
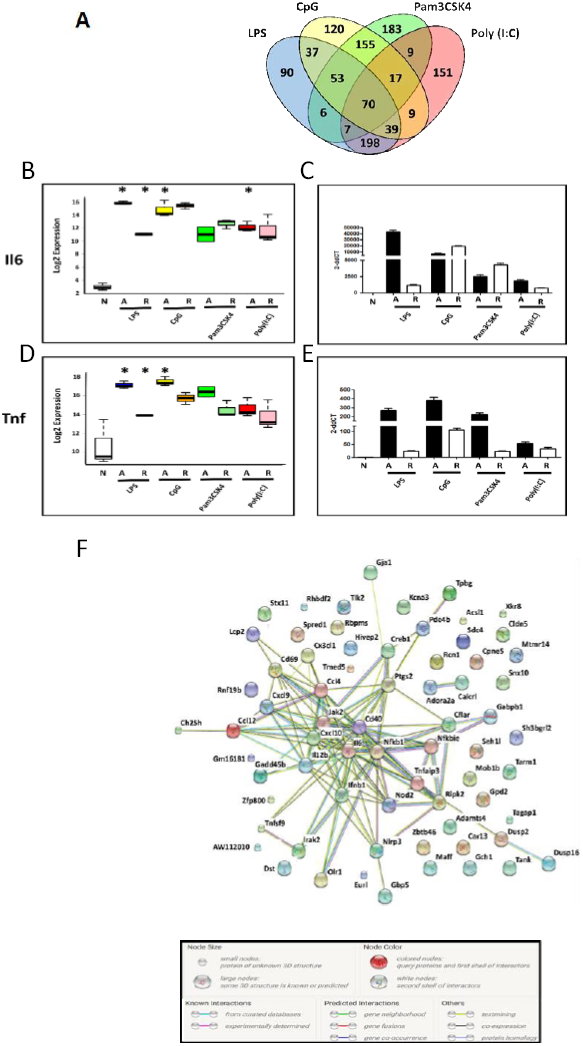
Acute activation of BMDM by TLR ligands. **(A)** Overlap of the top 500 genes induced at 4hrs by 100ng/ml LPS (blue), 1μM CpG (yellow), 100ng/ml Pam3CSK4 (green), 10μg/ml Poly (I:C) (red). **(B)** Microarray measurement (Log_2_) of *Il6* mRNA in naïve (N), Acute (A) or Re-stimulated (R) BMDM. **(C)** qRT-PCR of *Il6* mRNA. **(D)** Microarray (Log_2_) of *Tnf* mRNA. **(E)** qRT-PCR of *Tnf* mRNA. **(F)** StringDB protein-protein association network for the proteins encoded by the 70 genes commonly induced by the four TLR ligands indicated in Venn. Nodes: proteins. Edges: interactions from STRING database. For Box-whisker plots, median, min, max shown, n=2 or 3 samples * adjusted p < 0.05. For histograms, delta-delta CT normalised to normalised to naïve BMDM, n=3. See also Supplementary Figure 1: TF motif analysis of top 500 DE genes.

As 47% of the inducible gene set was ligand-specific, we searched for evidence of modules whose engagement was restricted to ligation of a single TLR. Clustering by Pearson correlation revealed a group of probes corresponding to 55 genes that were induced only in the acute Poly (I:C) condition (Figure 3). Enrichment of functional modules in this group of genes was largely driven by expression of *Fgf8* and *Ppm1a*, which are part of insulin, MAPK and Phosphoinositide 3-kinase (PI3K) signalling pathways. Additionally, genes in this group included candidates for anti-viral (*Papolb*) and immune signalling activities (*Ppm1a*, *Trim12c*, *Csmd2* and *Tbx21*). However, few of the 55 genes have been characterised extensively in an immune context, and 15% of this gene set consisted of genes uncharacterised in any context, highlighting the potential for further characterisation of the TLR3-induced transcriptome.

**Figure 3.**
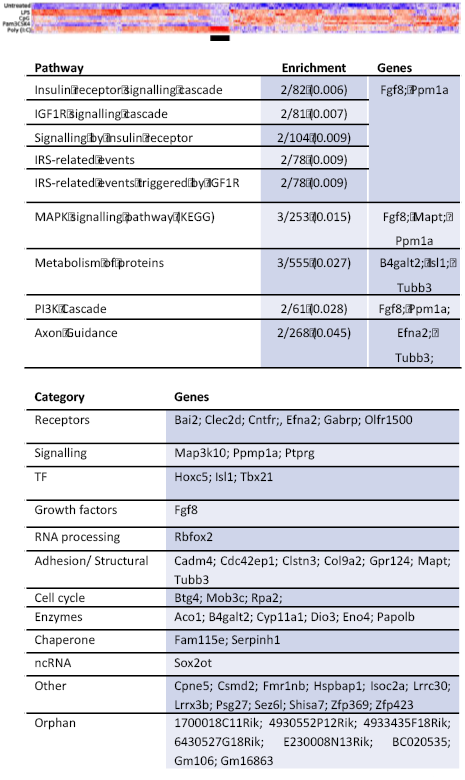
Characterisation of probes induced only by Poly (I:C) **(A)** Heat map (Pearson correlation) of combined list of top 500 acutely induced genes per acute condition. Colour indicates row z-score, ranging from −2 (dark blue) to 2 (dark red). Black bar indicates cluster of probes highly induced only in Poly (I:C). **(B)** Table of Reactome pathways enriched (adjusted p < 0.05) in genes induced uniquely by acute infection with Poly (I:C). Enrichment: number of genes in test list/number of genes in pathway (adjusted p-value). **(C)** Table of genes grouped by biological category.

Tolerised genes were defined as those significantly upregulated at 4 hours acute stimulation, but demonstrating at least 1.5-fold lower activation on re-stimulation. A total of 1644 genes matched this pattern in one or more condition (Supplementary table S1). The largest group of tolerised genes was shared between LPS and Poly (I:C) (Figure 4, cluster 1); the smallest subset of tolerised genes were tolerised in all conditions except Poly (I:C) (Figure 4 cluster 2); a substantial subset of genes was only tolerised under LPS stimulation (Figure 4, cluster 3); and the remaining genes were tolerised in all conditions (Figure 4, cluster 4). The promoter regions of genes that were tolerised by LPS and Poly (I:C) (Cluster 1) were enriched for both NF-κB and IRF motifs, whereas the subset of genes that were exclusively not tolerised by Poly (I:C) (Cluster 2) lacked the IRF motif. These were not tolerised because they were not acutely upregulated in the Poly (I:C) condition. Likewise, the genes that were only tolerised by LPS (cluster 3) were acutely upregulated only in this condition. The promoter regions of this gene set were largely dominated by the presence of an IRF4 and zinc finger motif. NF-κB p65-Rel promoter motifs were common to the majority of tolerised clusters, an observation consistent with previous links between NF-κB and tolerance (Yan *et al.*, 2012).

**Figure 4:**
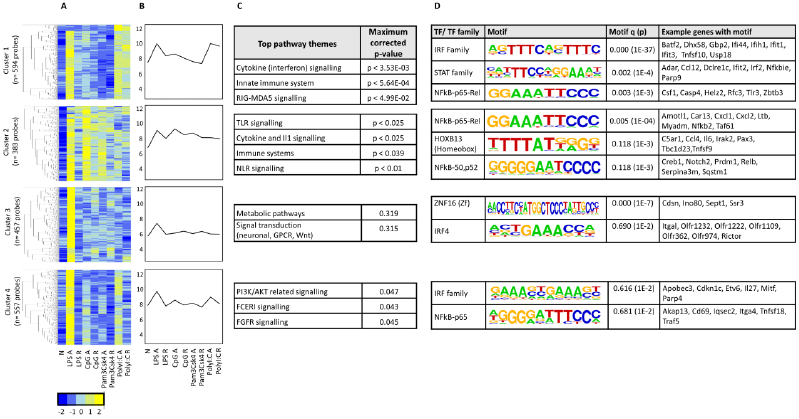
Tolerised probes demonstrate four patterns of expression. Probes showing tolerance in at least one infection were clustered using PAM. **(A)** Heat maps for each cluster. Colour indicates row z-score of mean log2 expression. X axis: treatment (N: Naïve, A: acute, R: re-infection). Y-axis: Pearson correlation of z scores. **(B)** Mean log2 expression pattern for all probes in each cluster (y-axis) per condition (x-axis). **(C)** Significantly enriched pathways, grouped thematically. Maximum adjusted p-value for all pathways significantly enriched in that theme are shown. **(D)** Transcription factor binding motifs enriched in each cluster. Motif logos and adjusted p-values are representative for each transcription factor family. Full motif enrichment results are available at www.stemformtatics.org.

These data revealed the surprising observation that the genes identified in any TLR-tolerance state are also always found in LPS-tolerance, indicating that LPS/TLR4-tolerance represents the archetype tolerisable state in BMDM. While LPS represented the most comprehensive tolerance condition, significant differences were observed in the capacity of other TLR to reduce pro-inflammatory cytokine and chemokine expression in the tolerant state. For example, while *Tnf* expression was consistently tolerised in all conditions, *Il6* was tolerised only in cells restimulated via LPS-TLR4 (see Figure 2). Similarly, *Ifnb1* was tolerised by ligands for TLR4 and TLR2, but not by TLR3 or TLR9, while *Il10* was tolerised by ligand for TLR3 and TLR4 but not TLR2 and TLR9 (Figure 5). IL-12 subunits were also differentially tolerised: *Il12a* was tolerised only in LPS conditions, whereas *Il12b* was tolerised by all conditions except Poly (I:C). These patterns are exemplified in Figure 5, and illustrated for TLR, NLR pathways, cytokines, growth factors and chemokines in Supplementary Figures S2-S4.

**Figure 5:**
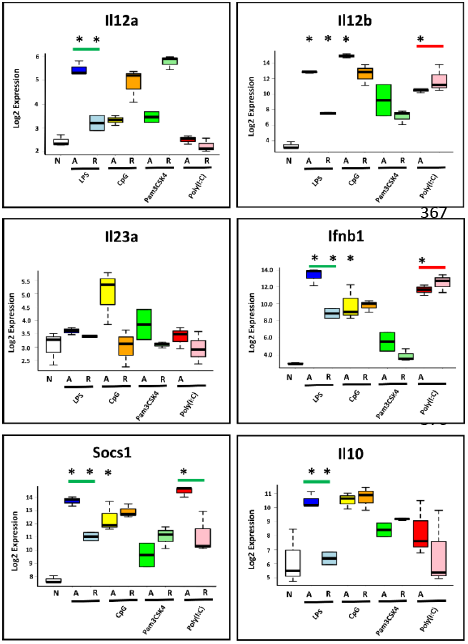
TLRs exhibit distinct patterns of tolerance. Microarray profiles of *Il12a*, *Il12b*, *Il23a*, *Ifnb1*, *Socs1*, *Il10*. Y-axis: Log2 expression. X-axis: conditions. See also Supplementary Figures S2-S4. Box and whisker plots are median, min, max. n=2-3.

### Innate memory – super-induction, super-repression and delayed responsiveness to TLR ligands

By definition acute induction was a pre-requisite of tolerance, however not all acutely-induced genes were tolerised. Arguably, tolerance represents one form of innate ‘priming’ or ‘training’, where pre-exposure to a PAMP alters macrophage responses to re-exposure. We identified 174 acutely induced genes that demonstrated further induction on re-stimulation with one or more TLR ligand (Figure 6, Supplementary table 1). Over a third of these genes were predicted to be secreted factors, including chemokines *Ccl2*, *Ccl5*, *Ccl8*, *Cxcl3*, *Cxcl5*; cytokines *Csf3*, *Ifnb1*, *Il1a*, *Il1b*, *Il6*, *Il12b*, *Il18bp*. As evident from Supplementary Figures S2-S4, many of these were tolerised or super-induced depending on the pathway of activation. Again, the dominant patterns were shared between TLR4/LPS and TLR3/Poly (I:C), with enrichment of IRF, bZip and NF-κB motifs in the promoters of these ‘super-induced’ genes.

**Figure 6:**
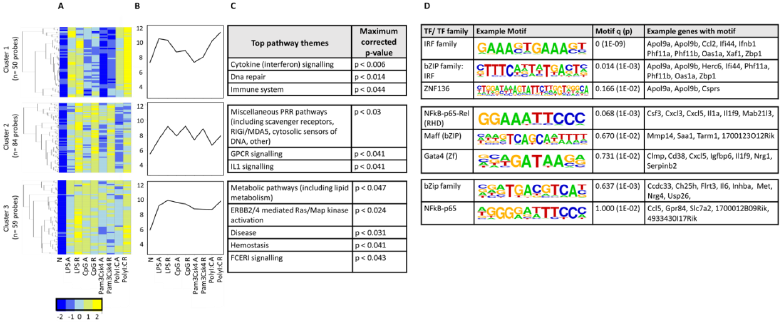
Ligand-specific patterns of super-induction on TLR re-stimulation. Probes showing super-induction in at least one infection were clustered using PAM. **(A)** Heat maps for each cluster. Colour indicates row z-score of mean log2 expression. X axis: treatment (N: Naïve, A: acute, R: re-infection). Y-axis: Pearson correlation of z scores. **(B)** Mean log2 expression pattern for all probes in each cluster (y-axis) per condition (x-axis). **(C)** Significantly enriched pathways, grouped thematically. Maximum adjusted p-value for all pathways significantly enriched in that theme are shown. **(D)** Transcription factor binding motifs enriched in each cluster. Motif logos and adjusted p-values are representative for each transcription factor family. Full motif enrichment results are available at www.stemformatics.org.

Genes that were transiently repressed, re-gaining at least 1.5-fold expression upon re-stimulation (Supplementary Figure S5, Supplementary table 1) were dominated by genes involved in metabolic respiration, with motif enrichment implicating ETS-factors as the major transcriptional regulator of this pattern. A smaller set of genes were acutely downregulated, then further strongly repressed on re-stimulation (Figure S6, supplementary table 1). These were predominantly genes implicated in cell cycle processes, and may reflect the culture system (mouse bone-marrow derived macrophages).

The set of 70 genes acutely induced in all conditions were overwhelmingly subjected to a tolerance pattern, with a small number exhibiting ligand-specific super-induction (Figure 7 (A)). This may indicate that coordinate mechanisms determine whether the expression of an inflammatory mediator is tolerised or trained. Indeed, the core members of NF-κB activation were themselves subjected to altered expression on re-stimulation with a TLR-ligand (Figure 7 (B)). Increased levels of the NF-κB p50 subunit and the IĸB protein BCL-3 were induced by all TLR ligands tested (Figure 7 (C)) underlying the previously established role for the p50:BCL-3 transcriptional repressor complex in limiting TLR responses (Carmody *et al.*, 2007). NF-κB target genes were highly represented in BMDM genes with a ‘memory’ status of the original stimulation, with the vast majority of these categorised in the tolerised group (Figure 7 (D)).

**Figure 7:**
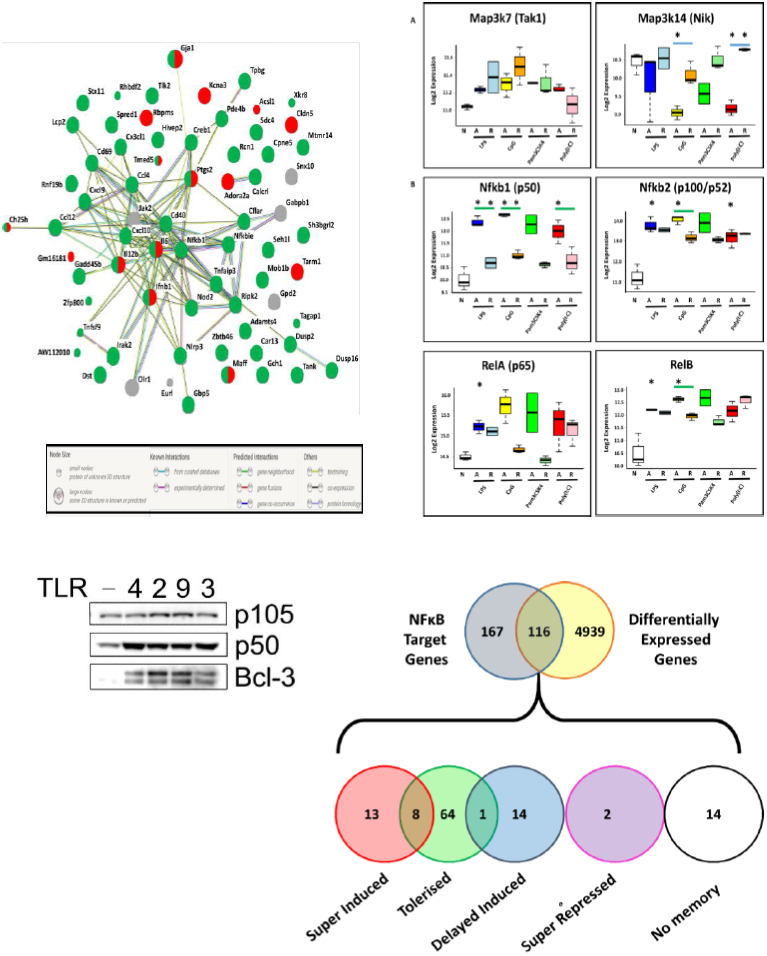
**(A)** StringDB protein-protein association network for the proteins encoded by the 70 genes commonly induced during acute responses by the four TLR ligands. Nodes: proteins. Edges protein-protein association. Colours: Green: tolerised; Red: super-induced; Grey: no memory pattern. **(B)** Key components of the canonical and non-canonical NF-κB pathways exhibit different regulatory patterns. Log2 expression of two kinases central to the canonical (left) and non-canonical (right) NF-κB pathways. Y-axis: Log2 expression. * indicates significant differential expression (adjusted p < 0.05). Coloured bars indicate memory patterns: green: tolerised; red: super-induced; magenta: super-repressed; blue: delayed-induced. **(C)** Western Blot analysis of p105, p50 and BCL3 expression in cells stimulated for 24 hours with ligand for TLR4, TLR2, TLR9 and TLR3 as indicated. **(D)** Characterisation of differential expression and memory status of NF-κB target genes. Overlap between NF-κB target genes and differentially expressed genes (adjusted p < 0.05, any acute infection). Number of genes per pattern are indicated in each coloured circle. Circle overlaps show numbers of genes with both patterns, either due to multiple differently behaving probes, or ligand-specific memory pattern differences.

## Discussion

Historically, the study of endotoxin tolerance was performed using lipopolysaccharides. Early research on endotoxin tolerance *in vivo* relied upon the fever response as a readout for responsiveness to endotoxin and led to the concept that tolerance was a hypo-responsive state due to a desensitisation to endotoxin. However, as the factors that mediate the innate inflammatory response were identified it became apparent that endotoxin tolerance is a state of altered responsiveness to stimulation rather than simply hypo-responsiveness. Initial transcriptomic analysis of LPS-induced tolerance in macrophage underscored this and revealed that a large number of LPS-inducible genes, particularly those encoding anti-microbial, anti-inflammatory and pro-resolution factors are not supressed during LPS tolerance (Foster, Hargreaves and Medzhitov, 2007). Remarkably, the same transcriptomic analysis has not previously been applied to tolerance induced by ligands for other TLRs and thus the relationship between tolerant states induced by specific TLRs remained unclear. In this study, we have addressed this by performing transcriptomic analysis of macrophages tolerised with ligands for TLR4, TLR2, TLR3 and TLR9. A comparative analysis of the transcriptomic profiles of TLR-specific tolerant cells provides us with fundamental insights into the molecular programming of the innate immune inflammatory response. Our data reveals that tolerance induced by each TLR is distinct and that TLR4 induced tolerance is the most comprehensive tolerant state relative to tolerance induced by other TLRs.

Our data reveals that the patterns of genes repressed during tolerance are largely associated with NF-κB dependent transcription regardless of TLR ligand, while IRF and B-ZIP motifs are overrepresented in the promoters of genes that are super-induced in tolerant cells. This likely reflects the pivotal role of the NF-κB transcription factor as a driver of pro-inflammatory gene downstream of all TLRs. Previous studies have established the importance of NF-κB in promoting inflammatory gene induction (Foster, Hargreaves and Medzhitov, 2007) and tolerance (Carmody *et al.*, 2007; Yan *et al.*, 2012) through the differential binding of NF-κB dimers to the promoters of repressed genes. The TLR-inducible expression of NF-κB target genes relies on the transactivation domains of NF-κB dimers containing a p65(RELA) or c-REL subunit. The transcriptional repression of NF-κB target genes during tolerance requires the binding of NF-κB p50 homodimers. The NF-κB p50 subunit lacks the transactivation domain found in the p65, c-REL and RELB subunits of NF-κB, and in the homodimeric form acts as a transcriptional repressor of NF-κB target genes. The stability of p50 homodimers is a key determinant of their repressor function and is controlled by polyubiquitination and proteasomal degradation. The IκB family member BCL-3 regulates p50 homodimer stability by inhibiting p50 ubiquitination and proteasomal degradation to form a stable DNA-bound repressor complex (Carmody *et al.*, 2007). Our data identifies elevated p50 and BCL-3 levels as a common feature of macrophages tolerised by all TLRs tested and suggests that this is a core mechanism for the repression of pro-inflammatory gene expression in TLR tolerant cells.

The repression of pro-inflammatory cytokine expression is one of the characteristic features of LPS tolerance. However, our analysis demonstrates that although each TLR ligand generally represses pro-inflammatory cytokine expression in tolerant cells, all cytokines are not universally tolerised and there is a highly diverse pattern of cytokine expression across all TLRs. *Tnf* is repressed in macrophages tolerised by all of the TLR ligands tested, however other important cytokines such as *Il6* are repressed in cells tolerised by TLR4 and TLR3 stimulation, but not by TLR2 and TLR9 activation. Of note, *Ifnb1* is repressed in cells tolerised by ligand for TLR4 and TLR2, but not by ligands for TLR3 or TLR9. The lack of repression of *Ifnb1* expression in cells tolerised by TLR3 ligand may reflect the importance of interferons in mediating an anti-viral immune response. This data suggests that sustained expression of *Ifnb1* in the context of a viral infection may be beneficial to host immunity. Similarly, the expression of the chemokines *Cxcl9* and *Cxcl10* by TLR3 tolerised macrophages correlates with the role of these factors in CD8+ T cell recruitment to sites of viral infection, cells that have a critical anti-viral activity (Thapa *et al.*, 2008). How these TLR-specific patterns of cytokine and chemokine expression are regulated is not known however the differential induction of negative regulators of TLR responses by individual TLRs may provide a potential mechanism. Thus, our data indicate specific programmes of TLR tolerance that are tailored towards the nature of the initiating stimulus. The immunological consequences of these specific patterns of cytokine repression will require further experimental investigation.

The differential profiles of cytokine expression in macrophages tolerised by different TLR ligands found in our analysis are also relevant to the more recently defined phenomenon of innate immune training. Innate immune training has been defined as enhanced innate host defence upon re-infection by the same or a different pathogen. Innate training is viewed as separate state to tolerance which is associated with the repression of inflammatory responses. However, our data suggests that this distinction may be too simplistic a categorisation of macrophage states following TLR activation. Our data shows that while *Tnf* expression is repressed in macrophages tolerised by all TLRs tested, *Il12a* shows an expression profile characteristic of training in cells tolerised by TLR9. Indeed our data is consistent with previous studies demonstrating a protective effect of CpG treatment against infection by *L. monocytogenes* that is accompanied by sustained IL-12 production (Krieg *et al.*, 1998). It is worth noting that innate training has been largely experientially defined by the enhanced expression of a limited number of cytokines. Our data caution against ascribing broad features of tolerance or training when studying TLR responses using a small number of experimental parameters. Rather, tolerance and training should be considered in a gene-specific context. This approach would likely better reflect the complex outcome of tolerisation of macrophage by individual TLRs as revealed by our analysis.

In summary, this study defines the transcriptional responses of macrophages tolerised with ligands for TLR2, TLR3, TLR4 and TLR9. Our data supports the concept that TLR tolerance promotes a shift away from a pro-inflammatory transcriptional response towards a response that is pro-resolution and anti-inflammatory in nature. The repression of transcription is generally associated with NF-κB target genes while genes with IRF motifs are more likely to be super-induced in tolerant cells. However, this study also reveals the differential repression of cytokines and chemokines in macrophages tolerised by specific TLR ligands. These patterns of expression may have functional relevance to stimulus specific inflammatory responses and may also be relevant to the study of innate immune training.

## Accessions

The data is available from (GSE81291) and Stemformatics (http://www.stemformatics.org/datasets/search?ds_id=6943)

## Acknowledgements

The authors thank Tyrone Chen and Othmar Korn and the Stemformatics team for microarray normalisation and visualisation. The authors acknowledge the support of the Medical Research Council (MR/M010694/1) (RC), and the Biotechnology and Biological Sciences Research Council (BB/M003671/1) (RC) and the COST Action BM1404 Mye-EUNITER (www.myeeuniter.eu), supported by COST (European Cooperation in Science and Technology), and the Australian Research Council (FT150100330; SR1101002) (CW). SB is funded by a scholarship from the University of Melbourne.

---

**Supplementary Figure 1:**
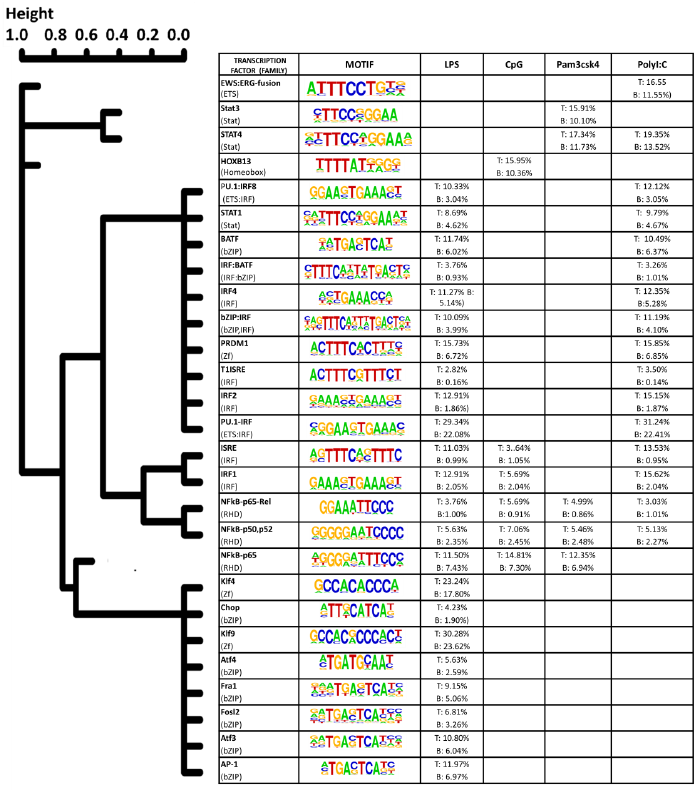
Transcription factor motif analysis of top 500 DE genes acutely regulated by TLR ligands in BMDM. Similarity tree grouping p-values across samples together. T: percent enrichment in test (DEG) set. B: percent enrichment in background set (background was set from all detected genes on microarray). Full transcription factor enrichment results are available at www.stemformatics.org.

**Supplementary Figures S2-S4:**
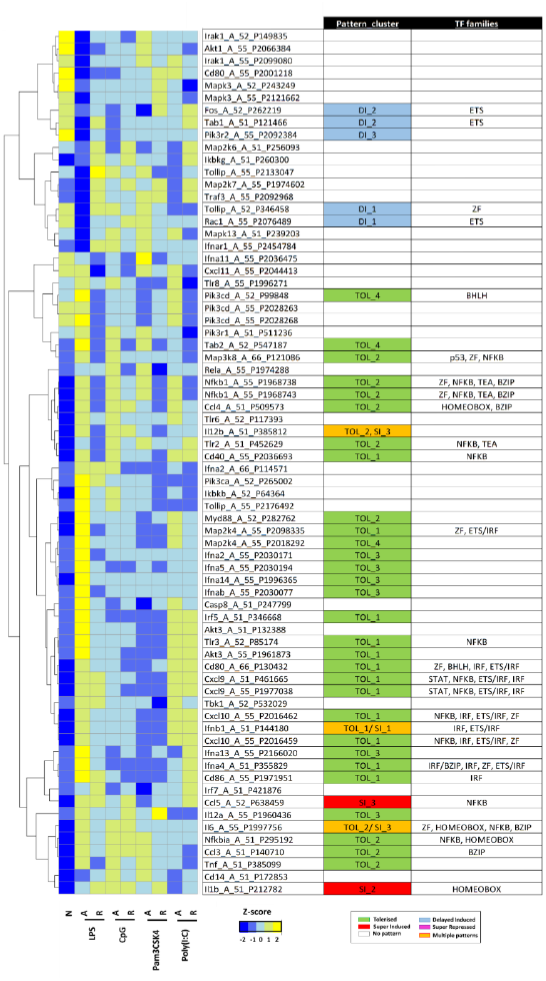
S2A Hierarchical clustering of TLR signalling pathway.

**S2B:**
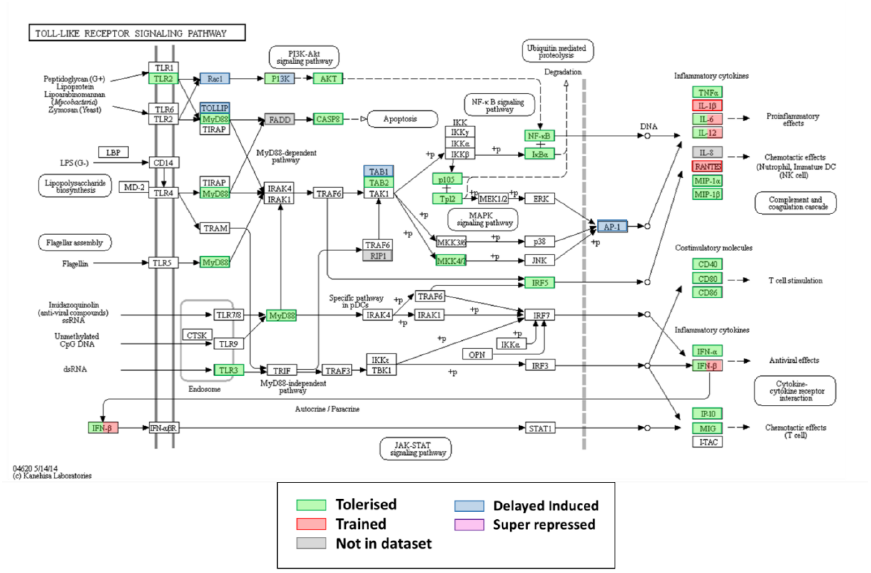

**S3A:**
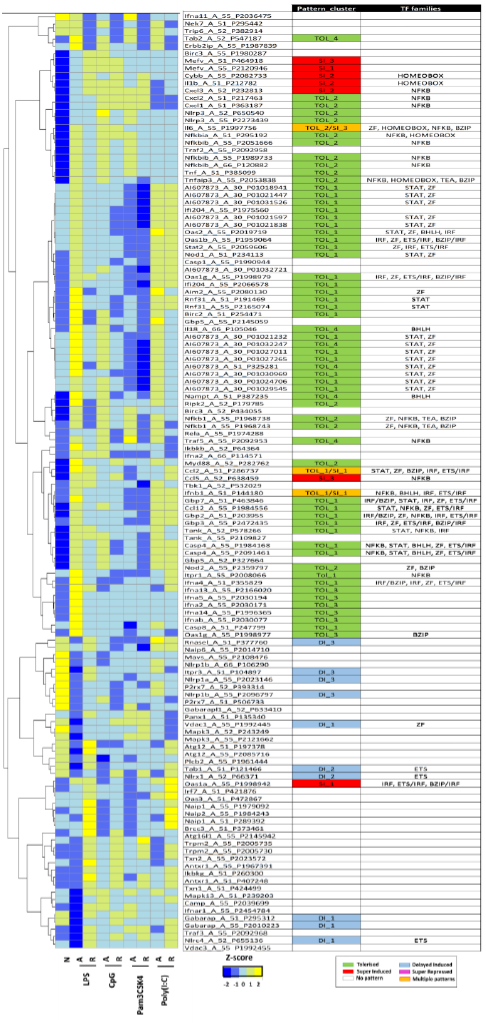
Hierarchical clustering of NLR pathway.

**S3B.**
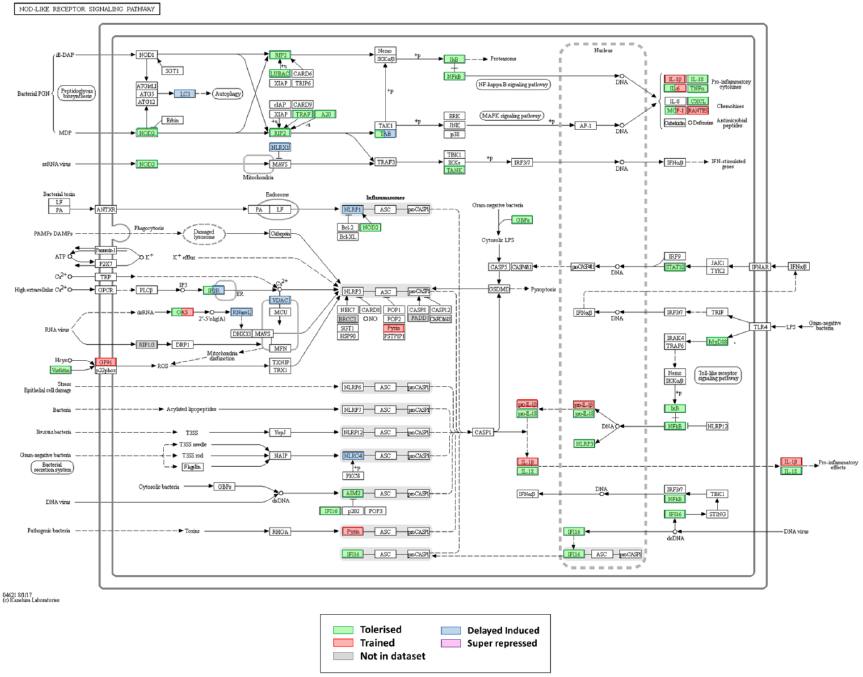

**S4:**
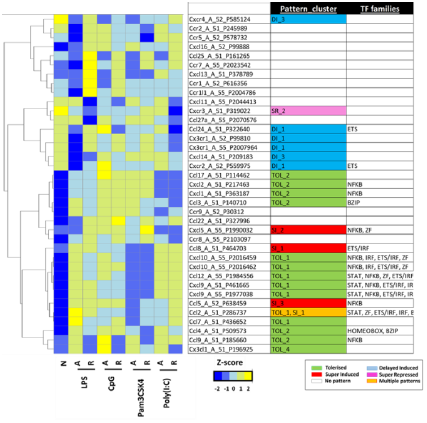
Hierarchical cluster of secreted factors and receptors, with chemokines highlighted.

**S4B.**
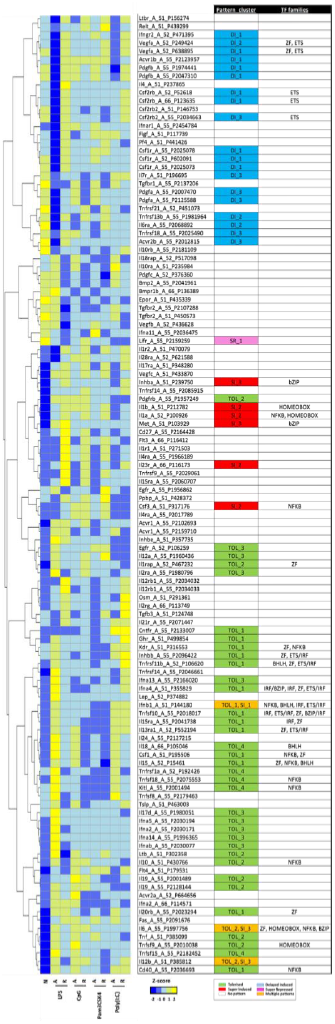
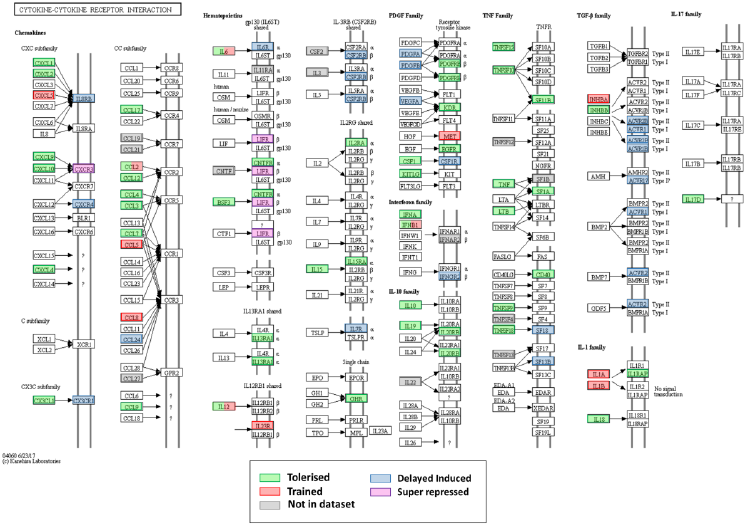

**Supplementary Figure S5:**
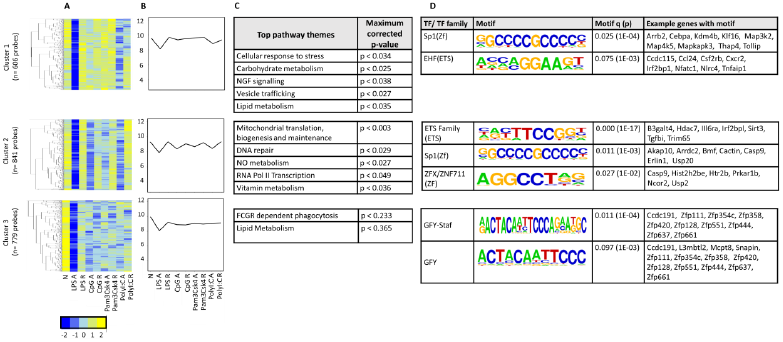
Probes that demonstrate transient repression. Probes showing transient repression in at least one infection were clustered using PAM. **(A)** Heat maps for each cluster. Colour indicates row z-score of mean log2 expression. X axis: treatment (N: Naïve, A: acute, R: re-infection). Y-axis: Pearson correlation of z scores. **(B)** Mean log2 expression pattern for all probes in each cluster (y-axis) per condition (x-axis). **(C)** Significantly enriched pathways, grouped thematically. Maximum adjusted p-value for all pathways significantly enriched in that theme are shown. **(D)** Transcription factor binding motifs enriched in each cluster. Motif logos and adjusted p-values are representative for each transcription factor family. Full transcription factor enrichment results are available at www.stemformatics.org.

**Supplementary Figure S6:**
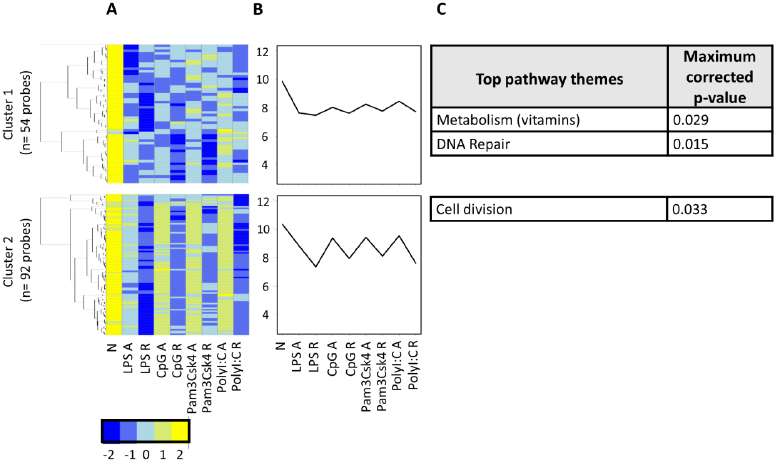
Probes that demonstrate consistent repression. Probes showing super-repression in at least one infection were clustered using PAM. **(A)** Heat maps for each cluster. Colour indicates row z-score of mean log2 expression. X axis: treatment (N: Naïve, A: acute, R: re-infection). Y-axis: Pearson correlation of z scores. **(B)** Mean log_2_ expression pattern for all probes in each cluster (y-axis) per condition (x-axis). **(C)** Significantly enriched pathways, grouped thematically. Maximum adjusted p-value for all pathways significantly enriched in that theme are shown.

**Table S2:** Top 5 GO terms (biological process) enriched for each set of TLR-ligand activated genes. Enrichment for each term is shown by the proportion of the GO list overlapping the TLR-induced gene lists (5), the number of terms in the TLR gene list /the total number of terms in the GO category. Significance of enrichment indicated by an adjusted p-value. Full list available in Table S1.

| LPS | CpG | Pam3csk4 | PolyI:C |
| --- | --- | --- | --- |
| **Innate immune response**<br>11% (87/771),<br>p = 7.95E-35 | **Innate immune response**<br>9% (71/771),<br>p = 8.36E-22 | **Innate immune response**<br>7% (55/771),<br>p = 7.03E-12 | **Innate immune response**<br>11% (81/771),<br>p = 2.37E-30 |
| **Defense response to virus**<br>19% (26/138),<br>p = 2.20E-14 | **Cellular response to LPS**<br>22% (20/93),<br>p = 1.27E-11 | **Cellular response to LPS**<br>16% (15/93),<br>p = 9.10E-07 | **Defense response to virus**<br>22% (32/138),<br>p = 2.88E-21 |
| **cellular response to IFN-beta**<br>59% (13/22),<br>p = 5.99E-14 | **inflammatory response**<br>12% (29/249),<br>p = 1.38E-10 | **inflammatory response**<br>9% (23/249),<br>p = 2.45E-06 | **cellular response to IFN-beta**<br>59% (13/22),<br>p = 4.55E-14 |
| **immune response**<br>12% (28/237),<br>p = 1.30E-10 | **immune response**<br>11% (27/237),<br>p = 1.29E-09 | **Response to LPS**<br>11% (18/162),<br>p = 5.67E-06 | **response to virus**<br>23% (21/90),<br>p = 1.08E-13 |
| **response to virus**<br>20% (18/90),<br>p = 2.55E-10 | **response to LPS**<br>14% (22/162),<br>p = 3.66E-09 | **Positive regulation of NF-kappaB activity**<br>14% (14/98),<br>p = 8.30E-06 | **negative regulation of viral replication**<br>46% (12/26),<br>p = 2.37E-11 |

